# Comparative and reproducible three-dimension functional spaces in primate cortical connectomes

**DOI:** 10.1101/2025.03.04.641091

**Authors:** Xiu-Xia Xing

## Abstract

Intrinsic activity or spontaneous oscillations are believed to be essential for the brain to maintain flexible, adaptive, and responsive cognition to both internal and external demands. However, a functional space remains elusive for understanding this inside-out organization in the primate cortex. Here, we leverage comparative fMRI analysis to identify three conserved and reproducible functional connectivity gradients as three axes of manifold (FMA), which span a low-dimensional organizational space of wakeful primate connectomes. In humans, the primary FMA covers a sensorimotor-to-transmodal gradient that supports signal detection for allostatic anticipation. The second FMA mirrors a representation-to-modulation gradient that integrates attention and performance monitoring to compute prediction errors. The third FMA profiles a self-to-goal gradient that links salience processing and episodic memory to update brain prediction by weighting errors and contextual relevance. Marmosets exhibit a homologous tripartite structure, suggesting evolutionary conservation of predictive coding motifs, albeit with species-specific topological variations. These findings unify predictive coding, allostasis, and intrinsic activity into a signal-error-salience predictive modeling framework, where sensory integration, error computation, and salience modulation interact to optimize adaptive responses. By bridging neurocomputational theory with empirical connectomics, this work establishes a cross-species blueprint of brain organization, offering experimental evidence or insights into conserved mechanisms of free-energy principles and dark-energy explorations. These FMAs highlight how evolution refines neural architectures while preserving core computational principles, paving the way for mechanistic studies of predictive dysfunction and evolutionary drivers of primate brain complexity.

---

Intrinsic oscillations are believed to be essential for the brain to maintain flexible, adaptive, and responsive cognition to both internal and external demands (*1*). The organization of such brain activity and its relationship to cognition and behavior have been increasingly approached through the lens of connectivity gradients (*2, 3*). This mathematical framework offers a way to analyze and represent complex neural data in spacetime, providing information on the underlying structures and dynamics of the organization of the primate brain (*4*). The concept of cortical gradients refers to continuous spatial variations in connectomic profiles across the cortex. It guides the brain to organize its connectivity along these gradients, with functional regions that smoothly transition from one state to another (*5*). By modeling these gradients as manifolds, researchers are now able to uncover how brain regions interact and how they change in response to different cognitive states or behaviors, with implications for linking brain geometry with function (*6*).

Previous research highlights how gradients in neuron density, connectivity, and gene expression shape primate cortical organization and suggests that gradients play a crucial role in brain function and offer insight into the evolution and development of brain networks between species (*7–9*). The cortical gradients derived using functional magnetic resonance (fMRI) data characterize the main axes of continuous variations in brain connectivity across the cortex. This forms a function manifold that reflects spatially organized patterns, where the cortical regions are hierarchically or topographically organized and related along the gradients. Such cortical gradient methods only keep a set of the strongest functional connectivity (e.g., the top 10% connectivity) for consideration (*10*). This threshold-based choice has derived consistent gradients of cortical organization between functional connectivity in the human brain and structural connectivity in the monkey brain (*5*). The tenet behind such a threshold choice is to approximate the underlying structural connectivity. However, the wave-like perspective of brain function or functionally traveling waves indicated that all connectivity are potentially significant properties of cortical wave propagation and can be biologically meaningful (*11–13*). Thus, a function manifold from the full fMRI connectivity data is essential to understand the cortical organization of primate brain function but remains elusive for modeling brain activity (*14*).

Comparative and reproducible analysis of function manifolds has been difficult due to the challenges of performing fMRI in wakeful non-human primate brains. In this work, we used two large-scale open datasets of resting-state fMRI (rfMRI) measurements from the Human Connectome Project (HCP) and the Chinese HCP (CHCP) to reconstruct the function manifold in the human cortical organization. Meanwhile, the same functional manifold analysis was also performed for wakeful marmoset brains based on two open rfMRI datasets from the National Institute of Health (NIH) and the Institute of Neuroscience (ION) at the Chinese Academy of Sciences.

## Materials and Methods

Wakeful resting-state fMRI data are from [Human Connectome Project](https://www.humanconnectome.org/) (HCP, N = 1003) (*15*) and the [Chinese HCP](https://www.scidb.cn/en/detail?dataSetId=f512d085f3d3452a9b14689e9997ca94&version=V4) (CHCP, N = 217) (*16*) for healthy human young adults while the NIH (N = 26) and the ION (N = 13) resource from the [Marmoset Brain Mapping Project](https://marmosetbrainmapping.org/) (*17*). All our analyses are based on group average preprocessed whole brain dense functional connectome (.dconn) data identifying temporal correlations between all cortical vertices and subcortical voxels. Specifically, each preprocessed individual data set was temporally trimmed and variance normalization applied and submitted to the principal component analysis (PCA) at the group level (MIGP) (*18*). The output of the group PCA with the spatial eigenvectors weighted in the upper N (N = 4500 for HCP, 2000 for CHCP, N = 1800 for NIH and ION) is then renormalized, reweighted and correlated to form the average dense connectomes at the group level (91, 282×91, 282 entries for the HCP/CHCP and 76, 457 × 76, 457 entries for the NIH/ION).

A neural manifold view of the brain indicates that brain function is commonly thought of as a low-dimensional space embedded in a very high-dimensional space where it is observed and measured (*19*). As illustrated in Figure 1, a 3D functional space is used to model the locally derivable manifold of highly complex and dimensional brain function, namely the functional manifold (FM). To estimate the FM’s axes (FMAs), diffusion map embedding was employed for identification of a set of low-dimensional manifolds (i.e., gradients) capturing principal dimensions of spatial variation in connectivity. In particular, this algorithm is implemented in BrainSpace toolbox (*10*) without thresholding functional connectivity to derive the functional affinity or similarity of the functional connectivity matrix. We note that all the connectivity (no threshold) are included in the computation for modeling the first three gradients span the ‘fully functional’ (not structurally approximated) manifold in the wakeful connectome organization.

**Figure 1:**
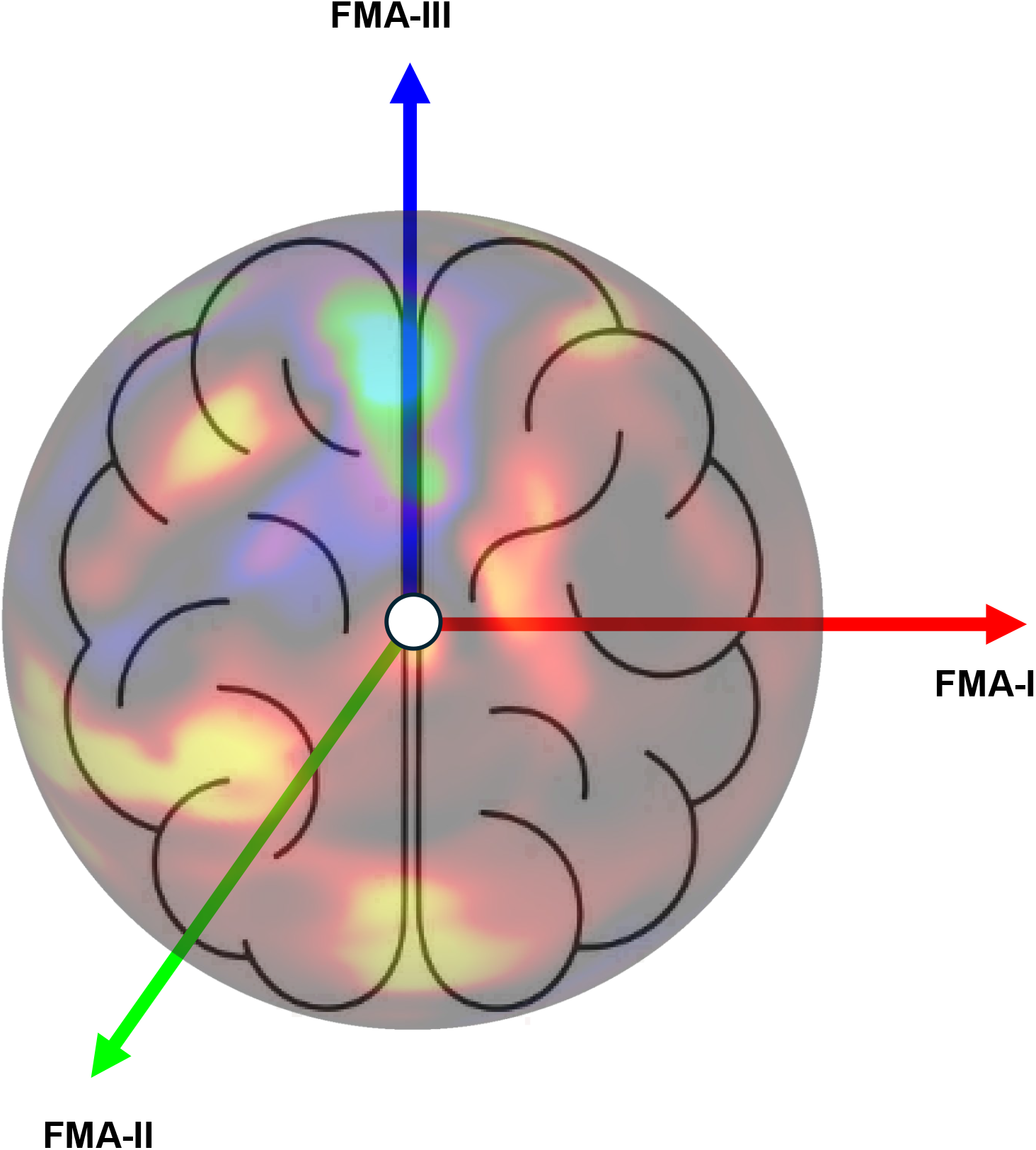
A neural manifold view of the primate brain function. Three inside-out axes of the functional manifold (FMA I-III) are the bases of a low-dimensional approximation of the high-dimensional functional space in the brain.

## Results

The Mega-ICA Group-PCA (MIGP) method (*18*) is used to construct a group-level connectome matrix from preprocessed rfMRI data, which are described in previous publications (*15–17*). It is designed to efficiently reduce the dimensionality of large-scale fMRI datasets while preserving meaningful functional connectivity patterns across individuals. MIGP generates dense connectomes from HCP, CHCP, NIH and ION datasets, respectively. These complete connectivity matrices are analyzed (that is, without the sparsity threshold) using BrainSpace toolbox (*10*), producing a series of gradients on the organization of the cortical connectivity. Specifically, the top three gradients are reproducible between independent datasets and explain a large portion of the observed variances in complete brain connectivity maps (HCP: 20.6%, 14.0%, 6.0%; CHCP: 20.3%, 12.2%, 6.6%; NIH: 46.8%, 19.2%, 6.5%; ION: 36.6%, 26.5%, 7.2%). We thus take the three gradients as a low-dimensional approximation of the high-dimension cortical connectivity space or its low-dimension geometry. From this perspective, these gradients form a three-dimensional manifold on the cortical connectivity geometry of the human brain function, that is, a function manifold. To visualize the functional manifold, we rescale each gradient to the same range from -1 to 1 and plot each vertex on the cortical surface as a point in the 3D space of the function manifold spanned by the three gradients as axes. As depicted in Figure 2 and Figure S1, the function manifold forms the surface of a regular tetrahedron or a functional representation in the human (Top Panel: HCP and CHCP) but not marmoset (Bottom Panel: NIH and ION).

**Figure 2:**
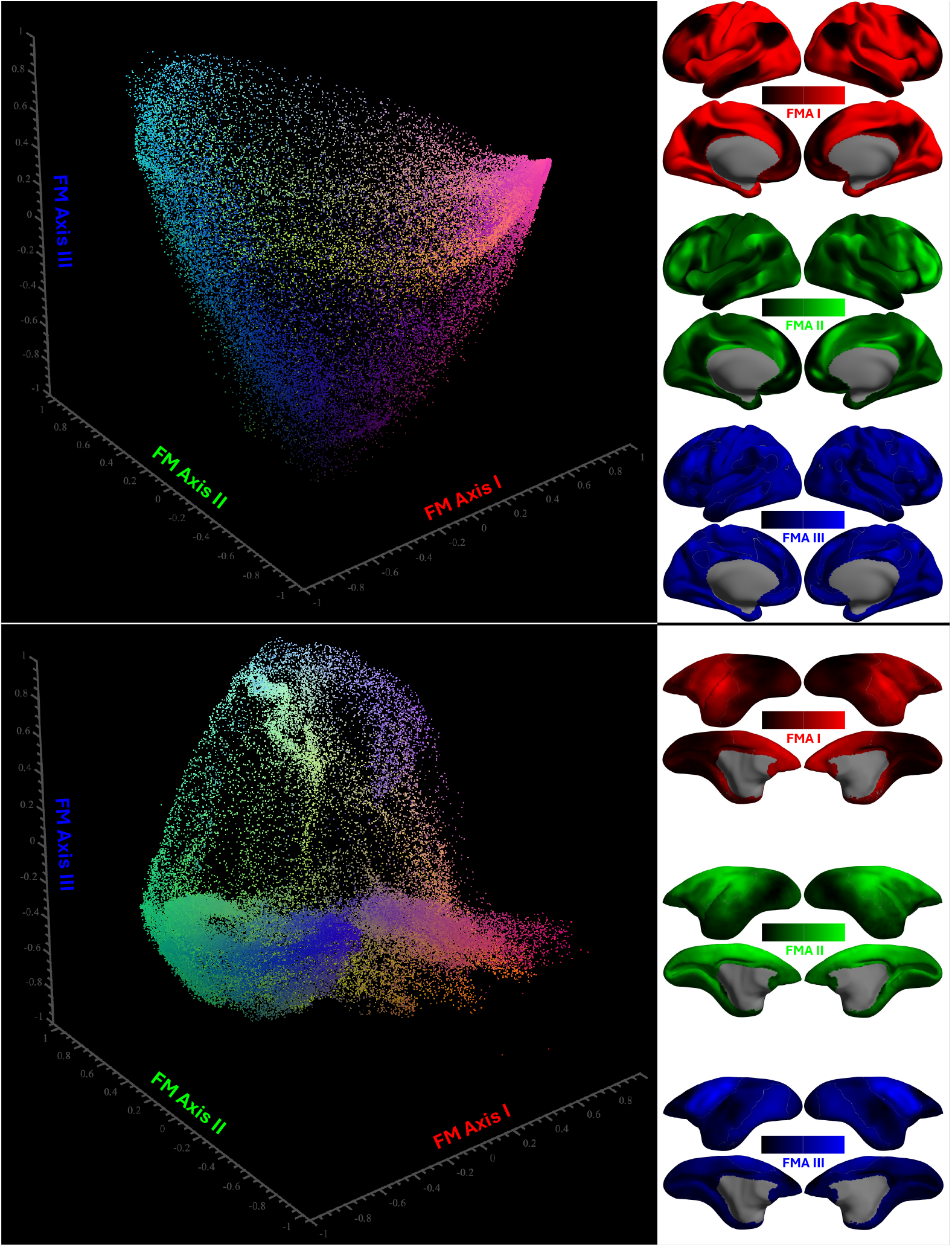
The 3D space of the functional manifold in primate cortical connectomes. Three axes are the three gradients of functional connectivity (gradient 1: red, gradient 2: green, gradient 3: blue). Given a vertex on the cortical surface, we plot a point in the 3D space with its coordinates as its three gradient values (Top panel: HCP, Bottom panel: NIH).

To further delineate anatomy of the function manifold in the human cortex, we extract mean rescaled gradient values of all the vertices within each of the 200 parcellating units (*20*) assigned to fifteen canonical cortical networks defined in DU15NET (*21*) for each hemisphere, and plot them along a spiral trajectory in ascending order (**gradient values are colored from the green-end inside to the yellow-end outside**). As shown in Figure 3A/B, the first FMA (FMA-I) starts inside the spiral from the frontoparietal network (FPN-B) and default-mode network (DMN-B and DMN-A), and ends outside the spiral at the premotor-posterior parietal rostral network (PM-PPr), action-mode network (AMN) (*22*), dorsal attention network (dATN-B), visual network (VIS-P). The second FMA (FMA-II, Fig. 1C/1D) starts inside the spiral from DMN-B and DMN-A, language network (LANG), and ends outside the spiral at FPN-A, salience and parietal memory network (SAL/PMN), dATN-A. The third FMA (FMA-III) starts inside the spiral from SAL/PMN, DMN-B, AMN, dATN-A, and ends outside the spiral at FPN-A and FPN-B, DMN-A (Fig. 3E/F). All the findings reported here are reproducible between HCP and CHCP (Fig. 3 and Fig. S2).

**Figure 3:**
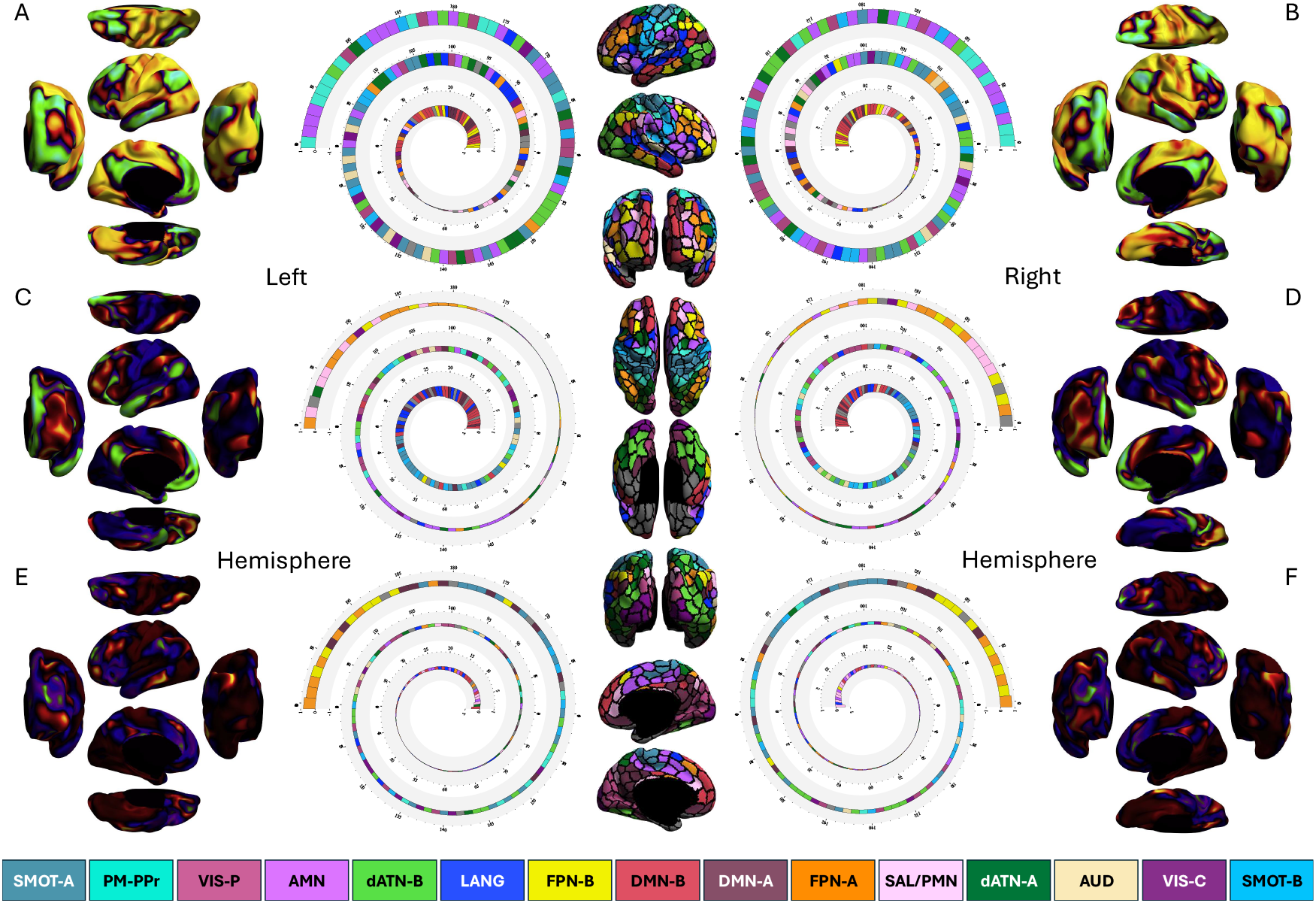
Top three gradient maps derived from the complete HCP connectivity matrix as axes of the functional manifold in the human cortical connectome. The left column (A,C,E) depicts the maps in the left hemisphere, and the right column (B,D,F) depicts the maps in the right hemisphere. The gradient values are averaged and ranked along spiral trajectories from the green-end inside to the yellow-end outside according to 200 parcellating units of each hemisphere (*20*). These parcels are colored and rendered onto the human cerebral cortex in terms of the fifteen canonical cortical networks defined in previous studies (*21,22*): Somatomotor-A (SMOT-A), Somatomotor-B (SMOT-B), Premotor-Posterior Parietal Rostral (PM-PPr), Action-Mode Network (AMN), Salience/Parietal Memory Network (SAL/PMN), Dorsal Attention-A (dATN-A), Dorsal Attention-B (dATN-B), Frontoparietal Network-A (FPN-A), Frontoparietal Network-B (FPN-B), Default-Mode Network-A (DMN-A), Default-Mode Network-B (DMN-B), Language (LANG), Visual Central (VIS-C), Visual Peripheral (VIS-P), and Auditory (AUD).

We delineate the anatomy of the function manifold in the marmoset cortex in a similar vein by extracting mean rescaled gradient values of all vertices within each of the 96 parcellating units assigned to fifteen canonical cortical networks defined in (*17*). As illustrated in Figure 4A/B, FMA-I starts inside the spiral from the high-level 2 visual-related network (highVN2), frontoparietal-like network (FPN), lateral primary visual and MT/MST visual-related network (VIS-l), default-mode-like network (DMN), and ends outside the spiral at the auditory and insular cortex salience-related network (AUD), premotor network (preMOT). FMA-II (Fig. 4C/D) starts inside the spiral from the high-level 1 visual-related network (highVN1), highVN2, orbital frontal network (orbFO), and ends outside the spiral at the dorsal somatomotor network (SMOT-d), FPN. FMA-III (Fig. 4E/2F) starts inside the spiral from the frontal pole network (FPO), preMOT, high-level 3 visual-related network (highVN3), AUD, and ends outside the spiral at the dorsal and ventral somatomotor networks (SMOT-d and SMOT-v). All the findings reported here are reproducible between NIH and ION (see Figure 4 and Figure S3).

**Figure 4:**
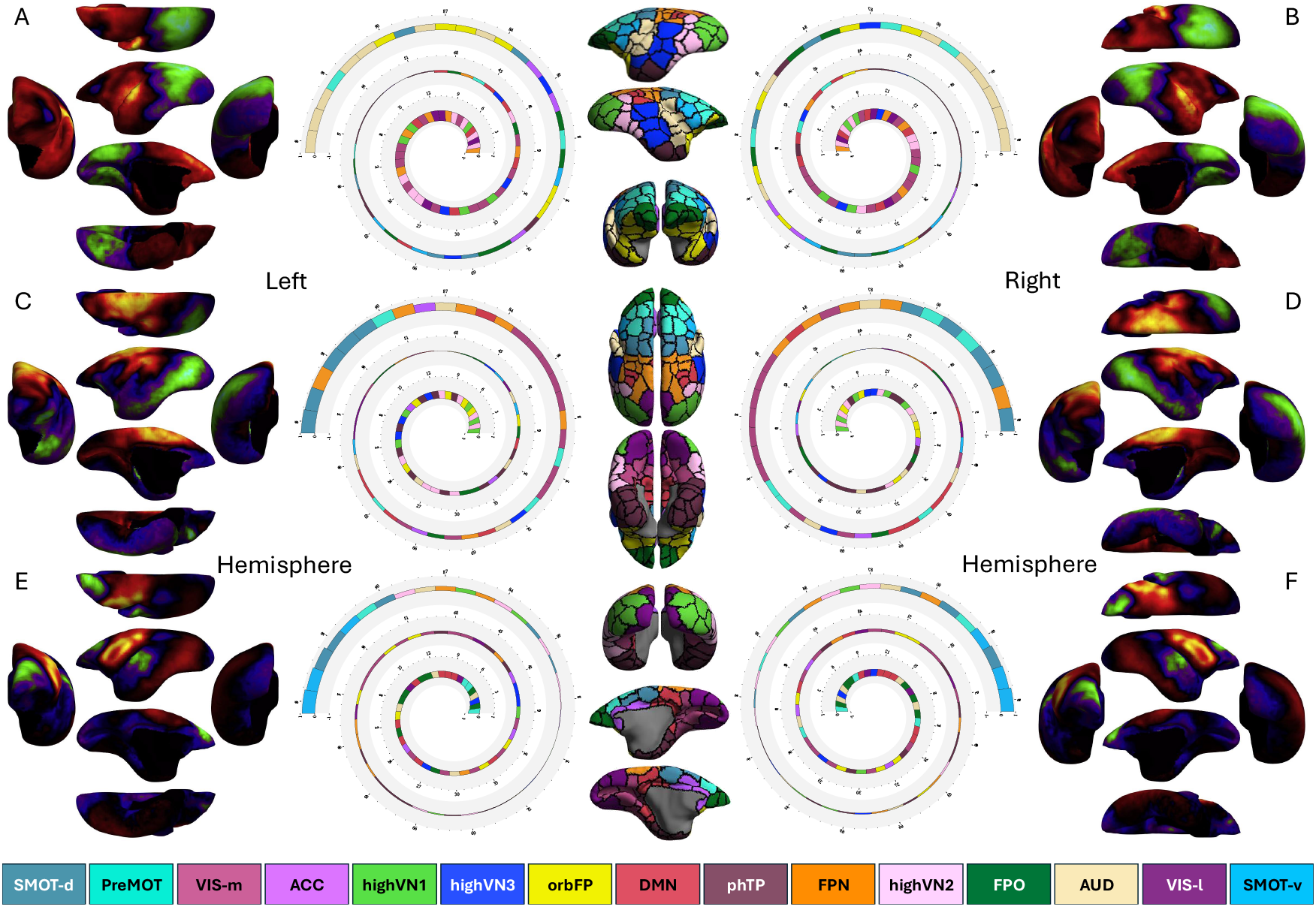
Top three gradient maps derived from the complete NIH connectivity matrix as axes of the functional manifold in the marmoset cortical connectome. The left column (A,C,E) depicts the maps in the left hemisphere, and the right column (B,D,F) depicts the maps in the right hemisphere. The gradient values are averaged and ranked along spiral trajectories from the green-end inside to the yellow-end outside according to 96 parcels in each hemisphere (*17*). These parcels are rendered onto the marmoset cerebral cortex in terms of the fifteen canonical cortical networks defined in (*17*) using the same colorset as in the human cortical network parcellation: dorsal somatomotor (SMOT-d), ventral somatomotor (SMOT-v), premotor (PreMOT), medial primary visual and MT/MST (VIS-l), lateral primary visual and MT/MST (VIS-m), auditory and insular cortex (AUD), anterior cingulate cortex (ACC), frontal pole (FPO), high-level 1 visual-related network (highVN1), high-level 2 visual-related network (highVN2), high-level 3 visual-related network (highVN3), orbital frontal (orbFP), frontoparietal-like network (FPN), default-mode-like network (DMN), the parahippocampus/temporal pole (phTP).

## Discussion

The brain operates as a highly dynamic and self-organizing system governed by the principles of free energy and predictive coding, where it continuously minimizes prediction errors through the constant comparison of sensory input with internal models (*23*). This enables the brain to anticipate and optimize responses to the environment, supporting allostasis, the regulation of internal states for adaptive survival (*24*). Meanwhile, the brain exhibits the feature of the cost of “dark energy” through spontaneous intrinsic neural activity (*1, 25*) to maintain homeostasis, prepare for future events and facilitate communication between functional networks, and supports efficient cognitive processing (*21*). These processes, from inside-outside interactions between the brain and the body to top-down predictions and bottom-up sensory integration, form a unified system that allows the brain to remain flexible, adaptive, and responsive to both internal and external demands. Our findings, for the first time, provide three reproducible dimensions of functional space in such a complex system as a basis to characterize its inside-out coordination observed by using fMRI to mathematically model primate connectomes (*14*).

Although the three axes (that is, gradient maps) of this functional space conceptually represent distinct aspects of characterizing the inside-out organization in the primate cortex (*26*), it is quite open and challenging to determine or interpret their exact roles in achieving predictive models of brain function. In human cortex, FMA-I has been documented as the sensorimotor-association, unimodal-transmodal or external-internal gradient for a core feature of the human brain organization to support a ‘domain-general’ motif (*5, 27*). We argue that it serves as a signal-detection motif to capture the predictable information flow along the external environment and the internal environment to serve the prediction of allostasis (*28*). FMA-II has a similar spatial profile to the ‘multiple demand system’ or the ‘representation-modulation system’ (*27*), forming an error-calculation motif to compute predictive errors by combining attention regulation, goal maintenance, strategy selection, performance monitoring with mental content. FMA-III has rarely been reported by previous gradient studies, although several recent work demonstrated some sparse evidence linking it to the recognition process of goal-directed and self-directed functions related to salience learning and episodic memory (*29–31*). We speculate on this axis as a salience-generation motif to support updates of the prediction by providing weights on both prediction signals and errors. In the marmoset cortex, we note that a similar structure of the inside-out functional space was observed. We argue that the three axes revealed form a signal-error-salience predictive model in marmosets may similar to humans but with a different shape as in Figures 2 and S1. However, understanding their comparative insights raises challenges and opportunities for future studies (*32*) because fMRI studies in awakened animals are much limited compared to those in humans (*33, 34*).

Our work produces comparative and reproducible low-dimensional representations of primate functional connectomes at the group level. This addresses functional references in the brain connectivity space for individualized FMA mapping methods. The spacetime dual regression (*35*) on these references will achieve more reliable and valid FMA measurements at the individual level (*36*).

## Acknowledgments

The authors thank Dr. Cirong Liu from the Institute of Neuroscience, Chinese Academy of Sciences for valuable discussions during the early preparation of the manuscript and sharing the marmoset dataset, and the National Basic Science Data Center for informatic support.

## Author contributions

Conception and Design: X-X.X. Data Analysis: X-X.X. Initial Drafting of the Manuscript: X-X.X. Critical Review and Editing of the Manuscript: X-X.X.

## Data and materials availability

All codes and MIGP-derived data are deposited in the Chinese Color Nest Data Community (CCNDC: https://ccndc.scidb.cn/en) in the Science Data Bank (https://doi.org/10.57760/sciencedb.24589).

## Supplementary materials

Figures. S1 to S3

## Supplementary Materials for

**Figure S1:**
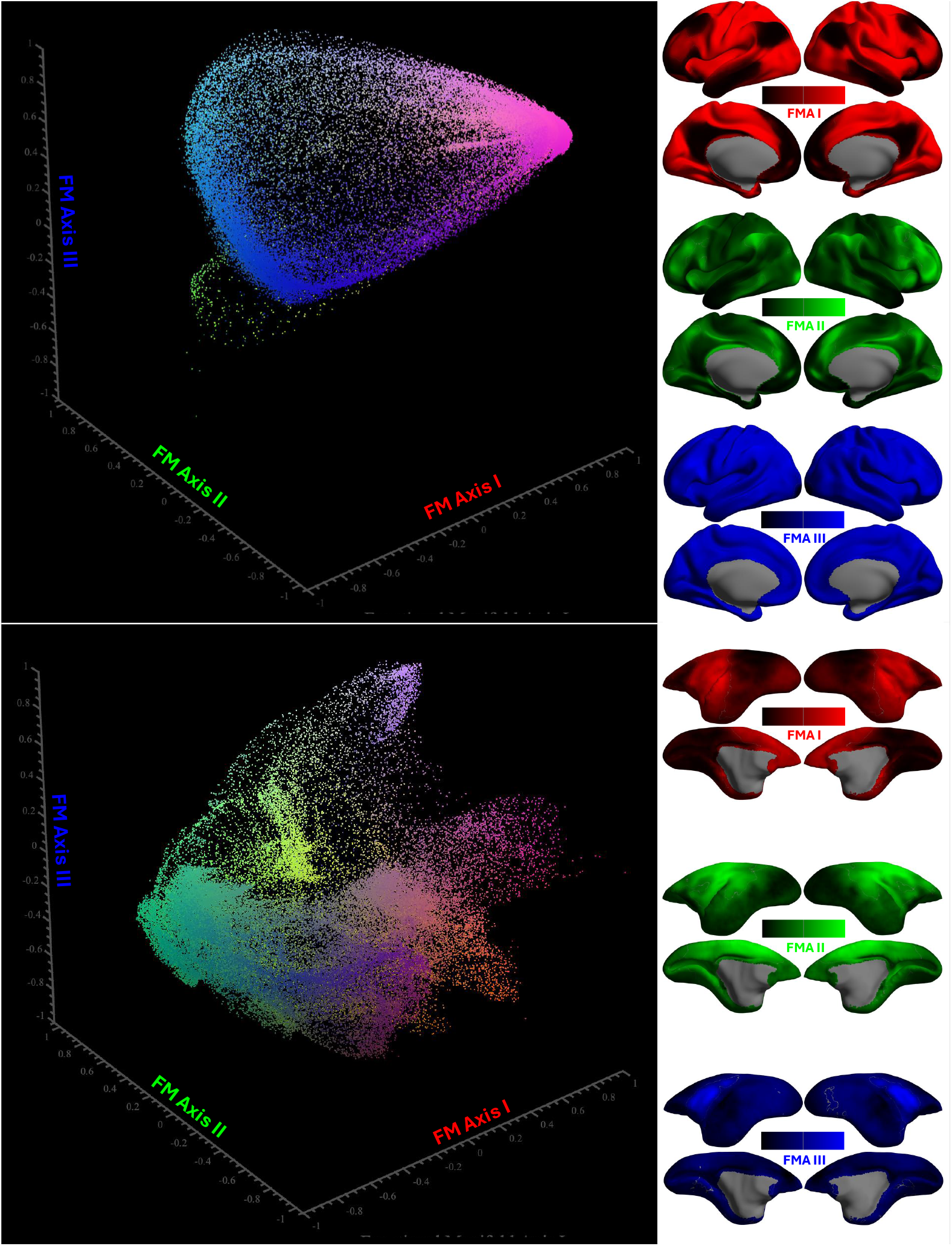
3D space of the functional manifold in primate cortical connectomes. Three axes are the three gradients of functional connectivity (gradient 1: red, gradient 2: green, gradient 3: blue). Given a vertex on the cortical surface, we plot a point in the 3D space with its coordinates as its three gradient values (Top panel: CHCP, Bottom panel: ION).

**Figure S2:**
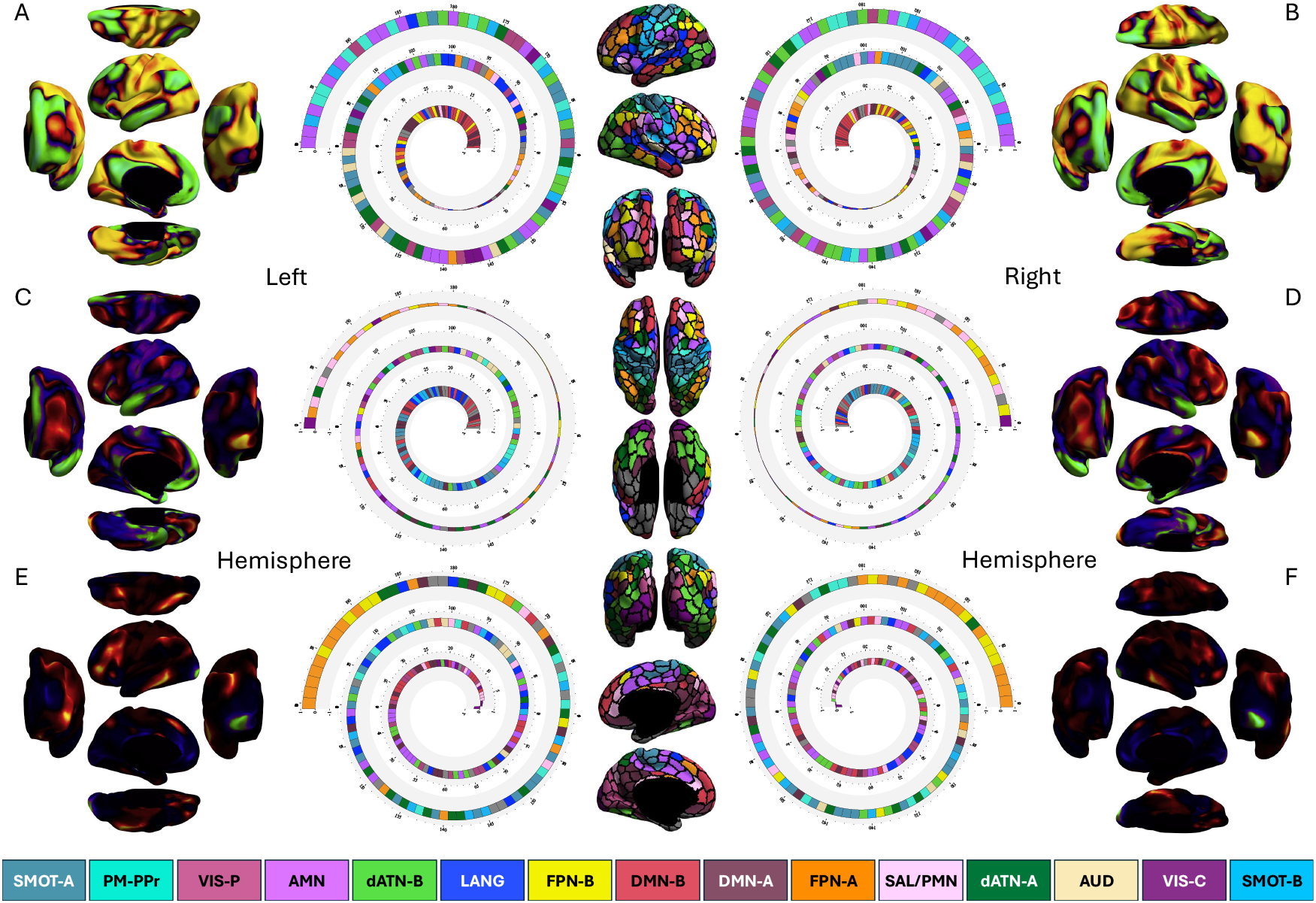
Top three gradient maps derived from the complete CHCP connectivity matrix as axes of the functional manifold in the human cortical connectome. The left column (A,C,E) depicts the maps in the left hemisphere, and the right column (B,D,F) depicts the maps in the right hemisphere. The gradient values are averaged and ranked along spiral trajectories from the green-end inside to the yellow-end outside according to 200 parcellating units of each hemisphere (*20*). These parcels are colored and rendered onto the human cerebral cortex in terms of the fifteen canonical cortical networks defined in previous studies (*21,22*): Somatomotor-A (SMOT-A), Somatomotor-B (SMOT-B), Premotor-Posterior Parietal Rostral (PM-PPr), Action-Mode Network (AMN), Salience/Parietal Memory Network (SAL/PMN), Dorsal Attention-A (dATN-A), Dorsal Attention-B (dATN-B), Frontoparietal Network-A (FPN-A), Frontoparietal Network-B (FPN-B), Default-Mode Network-A (DMN-A), Default-Mode Network-B (DMN-B), Language (LANG), Visual Central (VIS-C), Visual Peripheral (VIS-P), and Auditory (AUD).

**Figure S3:**
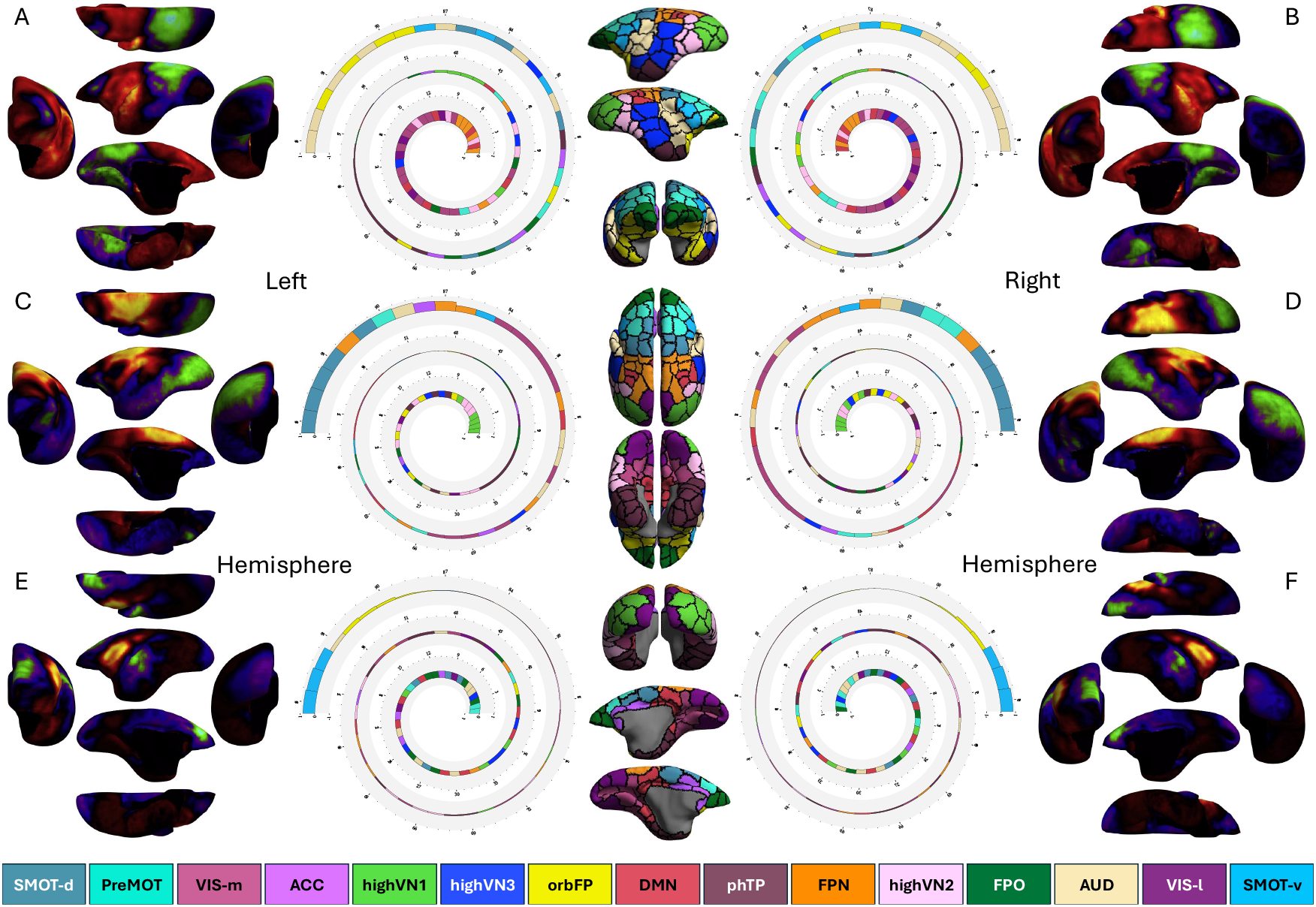
Top three gradient maps derived from the complete ION connectivity matrix as axes of the functional manifold in the marmoset cortical connectome. The left column (A,C,E) depicts the maps in the left hemisphere, and the right column (B,D,F) depicts the maps in the right hemisphere. The gradient values are averaged and ranked along spiral trajectories from the green-end inside to the yellow-end outside according to 96 parcels in each hemisphere (*17*). These parcels are rendered onto the marmoset cerebral cortex in terms of the fifteen canonical cortical networks defined in (*17*) using the same colorset as in the human cortical network parcellation: dorsal somatomotor (SMOT-d), ventral somatomotor (SMOT-v), premotor (PreMOT), medial primary visual and MT/MST (VIS-l), lateral primary visual and MT/MST (VIS-m), auditory and insular cortex (AUD), anterior cingulate cortex (ACC), frontal pole (FPO), high-level 1 visual-related network (highVN1), high-level 2 visual-related network (highVN2), high-level 3 visual-related network (highVN3), orbital frontal (orbFP), frontoparietal-like network (FPN), default-mode-like network (DMN), the parahippocampus/temporal pole (phTP).

